# Using compartmental models to understand excitation-inhibition imbalance in epilepsy

**DOI:** 10.1101/2023.11.03.565450

**Authors:** Aravind Kumar Kamaraj, Matthew Parker Szuromi, Daniel Galvis, William Stacey, Anne C Skeldon, John Terry

## Abstract

Epileptic seizures are characterized by abnormal synchronous bursting of neurons. This is commonly attributed to an imbalance between excitatory and inhibitory neurotransmission. We introduce compartmental models from epidemiology to study this interaction between excitatory and inhibitory populations of neurons in the context of epilepsy. Neurons could either be bursting or susceptible, and the propagation of action potentials within the brain through the bursting of neurons is considered as an infection spreading through a population. We model the recruitment of neurons into bursting and their subsequent decay to susceptibility to be influenced by the proportion of excitatory and inhibitory neurons bursting, resulting in a two population Susceptible – Infected - Susceptible (SIS) model. This approach provides a tractable framework to inspect the mechanisms behind seizure generation and termination. Considering the excitatory neurotransmission as an epidemic spreading through the neuronal population and the inhibitory neurotransmission as a competing epidemic that stops the spread of excitation, we establish the conditions for a seizure-like state to be stable. Subsequently, we show how an activity-dependent dysfunction of inhibitory mechanisms such as impaired GABAergic inhibition or inhibitory–inhibitory interactions could result in a seizure even when the above conditions are not satisfied.

## I. INTRODUCTION

Epilepsy is a complex neurological disorder affecting over 65 million people globally. Characterised by recurrent, unprovoked, seizures - the spread of pathological electrical activity in the brain - epilepsy can significantly impact both quality and quantity of life [1]. Seizures manifest in a variety of physical and psychological symptoms that are attributed to abnormal bursting of neurons in the brain. A common hypothesis for this abnormal bursting is an imbalance between excitatory and inhibitory neurotransmission which leads to increased levels of cortical excitability at the macroscale [2–4]. However, the precise mechanisms by which epileptic seizures begin and end are not well understood [5]. Experimental studies have shown that both altered excitatory [6, 7] and inhibitory [8–10] mechanisms can give rise to seizures.

From a theoretical perspective, most of the historic modelling effort in epilepsy has been devoted to computational models that accurately reproduce the features seen in electroencephalographic (EEG) data during seizures, normal brain activity and the transition between the two states [11–14]. Some of these models include dysfunctional inhibitory processes such as impaired GABAergic inhibition [15] and depolarising effects of GABAergic neurotransmission [16] to capture how epileptiform activity evolves over time under such conditions. As such, these models are valuable in explaining the epileptiform activity as seen in the EEG.

More recently, the focus of the modelling studies has extended beyond purely the reproduction of features seen in the EEG, with a focus on understanding the combined effects of dynamic and network mechanisms on the emergence of seizures [17]. For instance, a two-level Kuramoto framework has been used to distinguish between the brain networks constructed from the EEG of healthy controls and patients with epilepsy [18]. In another instance, a network model consisting of excitatory and inhibitory neurons has been applied to study the avalanche effect in neural networks and the factors that lead the network into an epileptic regime [19]. A simple phenomenological model based on codimension-one bifurcations has been used to classify epileptic seizures into several dynamotypes based on the signatures of critical transitions occurring at the seizure onset and offset [20].

In this paper, we introduce an alternative viewpoint - a tractable framework that captures the proportion of neurons firing within a population at any given time. Taking inspiration from compartmental models in epidemiology [21], we apply these in the context of epilepsy, using them to model the interactions between excitatory and inhibitory populations of neurons within a brain region. As such, we disconnect from providing a detailed physiological description of a single neuron or indeed a meanfield approximation of a population of neurons. Rather, our model allows for the selective activation and deactivation of neuronal populations and the subsequent study of their effects on the overall cortical circuitry. This is particularly important as experimental studies have shown that the role of excitation-inhibition imbalance in seizures may not be as simple as excessive excitation or reduced inhibition but rather the effects of this imbalance is both spatially [22] and temporally [23] contingent. This type of approach has been applied in a variety of contexts where dynamics spread across networks, including the spread of misinformation in social networks [24, 25], the spread of credit risk contagion in financial networks [26], and identifying the source of failure in power grid networks [27]. Modelling the propagation of electrical activity in the brain as an infection spreading through the neuronal population provides a suitable framework to study spatiotemporal effects through computer simulations.

Within this framework, neurons exist in either a bursting (i.e., “infected”) state or a quiescent (i.e., “susceptible”) state. We model the recruitment of neurons into bursting and their subsequent decay to susceptibility as influenced by the proportion of excitatory and inhibitory neurons bursting. This results in a two population Susceptible - Infected - Susceptible (SIS) model. The model is described by two first-order differential equations, making it possible to visualize the phase portrait of the system, including nullclines and equilibrium positions, and so enabling us to characterise the system dynamics. Our approach brings a fresh perspective to modelling candidate mechanisms underlying seizure generation and termination, through a simple, yet powerful framework.

The rest of the paper is arranged as follows. In Section II, we derive the equations governing the interaction between a single excitatory and inhibitory population of neurons based on compartmental models. Considering excitatory neurotransmission as an epidemic spreading through the neuronal population and the inhibitory neurotransmission as a competing epidemic that stops the spread of excitation, we establish the conditions for a seizure-like state to be stable. Subsequently, in Section III, we show how dysfunction of inhibitory mechanisms, such as impaired GABAergic inhibition and inhibitory–inhibitory interactions can create the environment for seizures to emerge. We discuss the implications of this modelling strategy and set out the future directions in Section IV. We conclude by summarizing our key observations in Section V.

## II. SIS MODEL IN EPILEPSY

One of the simplest compartmental models in epidemiology is the Susceptible-Infected-Susceptible (SIS) model. The model has two compartments or distinct states of the population - a susceptible (S) state and an infected (I) state. Susceptible individuals get infected upon exposure to infected individuals, with a rate of recruitment *β*. The infected individuals recover from the disease and return to susceptibility with a rate *γ*. The ratio of the two rates is called the basic reproduction number, 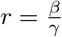 and measures the propensity of the disease to spread [28, 29].

In this section, we translate these ideas to epilepsy and establish what we call the ‘base model’. The base model has one excitatory and one inhibitory population and has equilibrium solutions descriptive of normal brain activity and a seizure.

To begin with, we consider neurons to exist in two states: a bursting or a tonic firing state, which we consider equivalent to the infected state in epidemiology, denoted by *B*, and a quiescent state, which we consider equivalent to the susceptible state, denoted by *S*. Mathematically, in most spiking neuron models, we have two attractors namely, a fixed point and a limit cycle [30, 31]. Physiologically, the fixed point attractor represents a quiescent state, which we model to be the compartment *S*. The limit cycle represents both bursting and tonic firing, which we model to be the compartment *B*.

Further, we consider two different populations of neurons, excitatory and inhibitory, denoted by subscripts *E* and *I* respectively. We formulate a compartmental model describing the interaction between these two populations of neurons according to the following rules:

- Bursting excitatory neurons (*B*_*E*_), upon interaction with susceptible excitatory neurons (*S*_*E*_), recruit them to burst. This can be thought of as analogous to how excitatory neurons cause other excitatory neurons to fire by sending a train of action potentials. The rate of recruitment is determined by the term *β*. Thus, 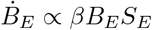.
- Bursting excitatory neurons (*B*_*E*_) decay to susceptible, proportional to the amount of neurons bursting at any given time. The rate of decay is defined by the term *γ*, and so is modelled as 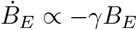. This term effectively describes a self-limiting mechanism for excitation: it is hard to sustain bursting when the proportion of neurons bursting is high as the neurons become depleted when firing for a long time.
- Bursting excitatory neurons (*B*_*E*_), upon interaction with susceptible inhibitory neurons (*S*_*I*_) recruit them to burst. This is designed to mimic how inhibition may work in the brain. When a higher proportion of excitatory neurons burst, a higher proportion of inhibitory neurons are activated to regulate their activity. The rate of recruitment is defined by the parameter *α*. Thus, 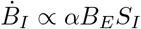.
- Bursting inhibitory neurons (*B*_*I*_) decay to susceptible determined by decay rate *ψ*. Thus, 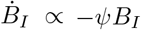.
- The total numbers of excitatory and inhibitory neurons in the system remain fixed, denoted by *N* and *M* respectively.

Following these rules, the equations governing the system can be expressed in dimensionless form as follows,

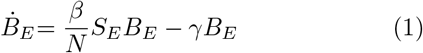

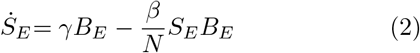

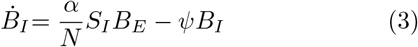

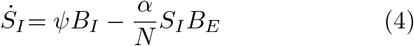

where *B*_*E*_ + *S*_*E*_ = *N* and *B*_*I*_ + *S*_*I*_ = *M* . Re-scaling the equations so that *B* and *S* represent the proportion of each population, i.e., *N* = *M* = 1, the system reduces to:

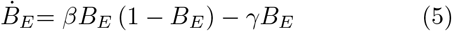

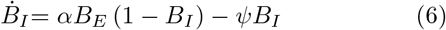

with dynamics restricted to Ω = [0, 1] × [0, 1]. In the above equations, we consider the terms *β* (*B*_*E*_, *B*_*I*_) and *γ* (*B*_*E*_, *B*_*I*_) to be functions of the current inhibitory and excitatory bursting populations. These functions represent how the proportion of neurons currently bursting affects the recruitment and decay of excitatory neurons respectively. The terms *α* and *ψ* are assumed to be constants in the base model.

This approach permits the study of interactions between an excitatory and an inhibitory population of neurons in a simple way, using two first-order ODEs. Because of this, it is possible to visualize the phase portrait of the system, including nullclines and equilibrium positions, for different choices of the functions *β* (*B*_*E*_, *B*_*I*_) and *γ* (*B*_*E*_, *B*_*I*_). This enables meaningful insights into system behaviour to be established.

### A. Nullclines and fixed points of the system

From Eq. (5), the *B*_*E*_ nullclines are given by *B*_*E*_ = 0 and 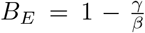. The second or the non-trivial arm of the *B*_*E*_ nullcline depends on the choice of the terms *β* (*B*_*E*_, *B*_*I*_) and *γ* (*B*_*E*_, *B*_*I*_). We describe this further in due course. The *B*_*I*_ nullcline can be derived from Eq. (6) as,

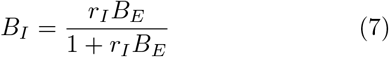

Where 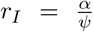 denotes the propagation ratio for inhibitory neurotransmission. The propagation ratio is analogous to the basic reproduction number for an infectious disease. In epidemiology, the basic number of reproduction is defined as the expected number of infections directly generated by one infected person in a population where all individuals are susceptible to infection [32]. We define two propagation ratios in our model, one for inhibitory and the other for excitatory neurotransmission. As bursting excitatory neurons are considered to recruit both inhibitory and excitatory neurons into bursting, the propagation ratio for inhibitory (excitatory) neurotransmission measures the net post-synaptic inhibitory (excitatory) activity generated due to a single bursting excitatory neuron assuming all the post-synaptic neurons are susceptible.

The equilibrium positions of the system are given by the points of intersection of the *B*_*E*_ and *B*_*I*_ nullclines. At (*B*_*E*_, *B*_*I*_) = (0, 0), the *B*_*E*_ = 0 arm of the *B*_*E*_ null-cline and the *B*_*I*_ nullcline intersect, giving rise to a fixed point. In epidemiology, this fixed point corresponds to the disease-free state. In the context of the brain, this represents a state where none of the neurons are firing and consequently, we will call this as the activity-free state. Depending on the choice of *β* (*B*_*E*_, *B*_*I*_) and *γ* (*B*_*E*_, *B*_*I*_), the second arm of the *B*_*E*_ nullcline and the *B*_*I*_ nullcline will intersect at one or more points, which epidemiologically reflect different endemic states. The fixed points could either be stable, unstable, or saddles. In the brain, these endemic states reflect steady state behaviour where a constant proportion of excitatory and inhibitory neurons are firing. We consider the stable endemic fixed points to be different manifestations of normal activity. Further, we consider the fixed point where the *B*_*E*_ coordinate is one (i.e., 100% of the excitatory neurons firing) to denote the seizure-like state. We use the collective term ‘functional brain states’ to represent the aforementioned fixed points in the context of the brain. A summary of various terms introduced so far in the paper is given in Table I.

Since our model is a simplified mathematical representation of neuronal activity, we have chosen the scenarios where no neurons are bursting and all excitatory neurons are bursting to serve as ideal representations of extreme physiological activity. Consequently, a realistic interpretation of the activity-free state is that physiologically, it corresponds to a state where the neuronal activity is so low that it ceases to sustain life. At the other end of the spectrum, a realistic interpretation of the seizure-like state would be that a vast majority of excitatory neurons are firing in unison, representative of an electrographic seizure.

### B. Modelling the interaction between the excitatory and inhibitory populations

We consider the interaction between a single excitatory and a single inhibitory population through appropriate choices of *β* (*B*_*E*_, *B*_*I*_) and *γ* (*B*_*E*_, *B*_*I*_).

To begin with, there is an inherent rate of recruitment of excitatory neurons into bursting and their subsequent decay into susceptibility, independent of the effects of inhibition. The inherent rate of recruitment should reach a maximum in the absence of inhibition whereas the inherent rate of decay should reach a minimum in the absence of inhibition.

Considering inhibition, it should inversely influence the bursting of excitatory neurons and proportionally increase the decay to susceptibility. Assuming both recruitment and decay vary linearly with respect to *B*_*I*_, we propose *β* (*B*_*E*_, *B*_*I*_) = *b* (1 − *B*_*I*_) and *γ* (*B*_*E*_, *B*_*I*_) = *c* (1 + *ϕB*_*I*_), where *b* and *c* are constants denoting the rates of recruitment and decay respectively. Similarly to the inhibitory population, the propagation ratio could be defined for the excitatory population as well, given by, 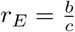.

*ϕ* scales the effect of inhibition on decay of bursting excitatory neurons relative to their inherent decay rate. A similar scaling term is not used to model recruitment since we require the recruitment to reach the maximum rate, *β*_max_ = *b* when *B*_*I*_ = 0, and reach zero when all inhibitory neurons are active, *β*_min_ = 0 when *B*_*I*_ = 1.

Substituting the proposed choices for *β* (*B*_*E*_, *B*_*I*_) and *γ* (*B*_*E*_, *B*_*I*_), the non-trivial arm of the *B*_*E*_ nullcline becomes,

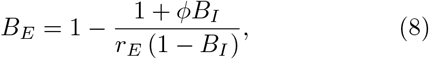

which exists on the domain of interest Ω only when *r*_*E*_ *>* 1.

When *r*_*E*_ *<* 1, only the *B*_*E*_ = 0 arm of the *B*_*E*_ nullcline exists. The only fixed point in this case would be, (*B*_*E*_, *B*_*I*_) = (0, 0), which is stable. This fixed point corresponds to the activity-free state, where there is no bursting activity in neurons. Consequently, to sustain propagation of activity within the neuronal population, *r*_*E*_ must always be greater than one.

When *r*_*E*_ *>* 1, the (0, 0) fixed point loses its stability. A second fixed point gets created at the intersection of the non-trivial arm of the *B*_*E*_ nullcline, given by Eq. (8), and the *B*_*I*_ nullcline. The coordinates of this fixed point are given by,

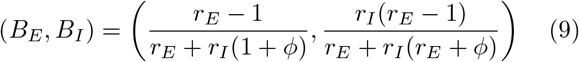

This fixed point corresponds to normal brain activity and is always stable when *r*_*E*_ *>* 1.

The phase portrait of the system, with the *B*_*E*_ and *B*_*I*_ nullclines represented as green and blue curves respectively, is shown in Fig. 1. The direction of the vector field as arrows and its magnitude as a colour map are indicated therein. From the phase portrait, it can be seen that the system has two fixed points: the activity-free state which is a saddle and indicated as a square, and the normal state which is stable and indicated as a circle. Furthermore, for initial conditions close to the x-axis (*B*_*I*_ ≈ 0, no inhibition), the direction of the vector field is determined by the value of the propagation ratio of excitatory neurotransmission, *r*_*E*_. When *r*_*E*_ is small, a relatively smaller proportion of excitatory neurons is recruited in comparison to the proportion of inhibitory neurons recruited, and the vector field points more towards up. When *r*_*E*_ is large, a relatively larger proportion of excitatory neurons is recruited in comparison to the proportion of inhibitory neurons recruited, and the vector field points more toward the right. Eventually, the system settles on to the normal state, which is a stable spiral. The rate of convergence drops down as the trajectories come closer and closer to the fixed point - this is readily seen in the magnitude of the vector field being low near the fixed point relative to elsewhere.

**FIG. 1.**
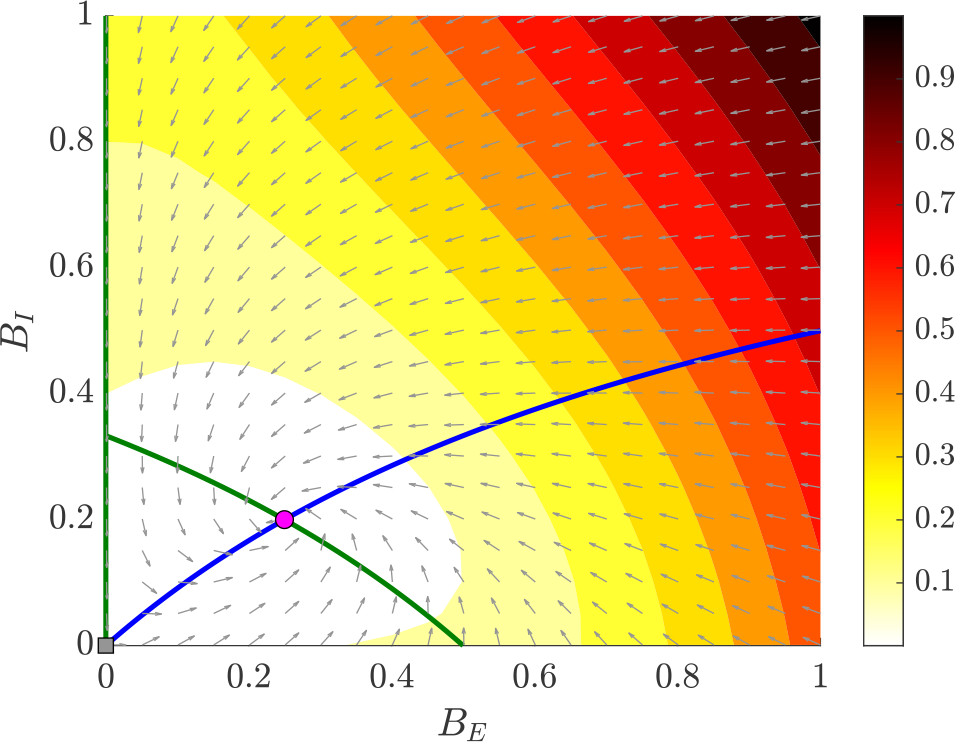
The phase portrait of the system for *β* = *b* (1 − *B*_*I*_) and *γ* = *c* (1 + *ϕB*_*I*_).

### C. Second order effects in bursting

As described in Section II, the decay of bursting neurons into susceptibility being proportional to the fraction of neurons currently bursting brings about a self-limiting effect. The physiological motivation for modelling this self-limiting behaviour in bursting comes from the refractoriness of neurons: it is hard to sustain bursting when the proportion of neurons bursting is high, since neurons become depleted after firing for a long time. Consequently, this self-limiting effect stabilises the system against perpetual bursting. However, to model an active seizure-like state in the system, interactions that counteract this self-limiting behaviour need to be considered.

The relative refractory period of neurons holds the key to introducing these competing effects. During the relative refractory period, the neurons are in a hyperpolarized state. Hence, a relatively higher stimulus is required to cause the neuron to fire. When a significant proportion of neurons in a population are bursting at a given time, it is reasonable to assume that a high level of activity could provide an increased stimulus and offset the effects of relative refractoriness. Hence, self-interaction of excitatory neurons could help sustain the bursting activity. Mathematically we model this sustenance in bursting activity using a second order interaction, 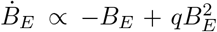 where the negative *B*_*E*_ term represents the original decay and the 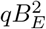 term serves to offset its effect. We call *q* ∈ [0, 1] the sustenance parameter since it quantifies the magnitude of bursting activity sustained due to second-order interactions.

Accordingly, we now consider *γ* = *c* (1 + *ϕB*_*I*_) (1 − *qB*_*E*_) to model the second-order effects in excitatory neurotransmission. The net decay in the bursting activity, *γB*_*E*_ = *c* (1 + *ϕB*_*I*_) *B*_*E*_ (1 − *qB*_*E*_) as a function of the sustenance parameter *q* is shown in

Fig 2. It can be seen that for *q* ≤ 0.5, it is progressively less challenging to sustain bursting, but it does not become easier since the net decay *γB*_*E*_ is monotonically increasing with decreasing slope. If 0.5 *< q <* 1, then beyond a critical value of *B*_*E*_, the slope of *γB*_*E*_ changes from positive to negative. Beyond this critical value, it becomes progressively easier to sustain bursting as more and more excitatory neurons are recruited into bursting, creating a positive feedback loop. When *q* = 1, the sustenance counterbalances decay at *B*_*E*_ = 1, leading to no excitatory neurons being pulled out of bursting (*γB*_*E*_ = 0), as shown in Fig 2. This counterbalance arising from setting *q* = 1 has important consequences, that we consider in Section II D. For now, we will examine the case 0 *< q <* 1.

**FIG. 2.**
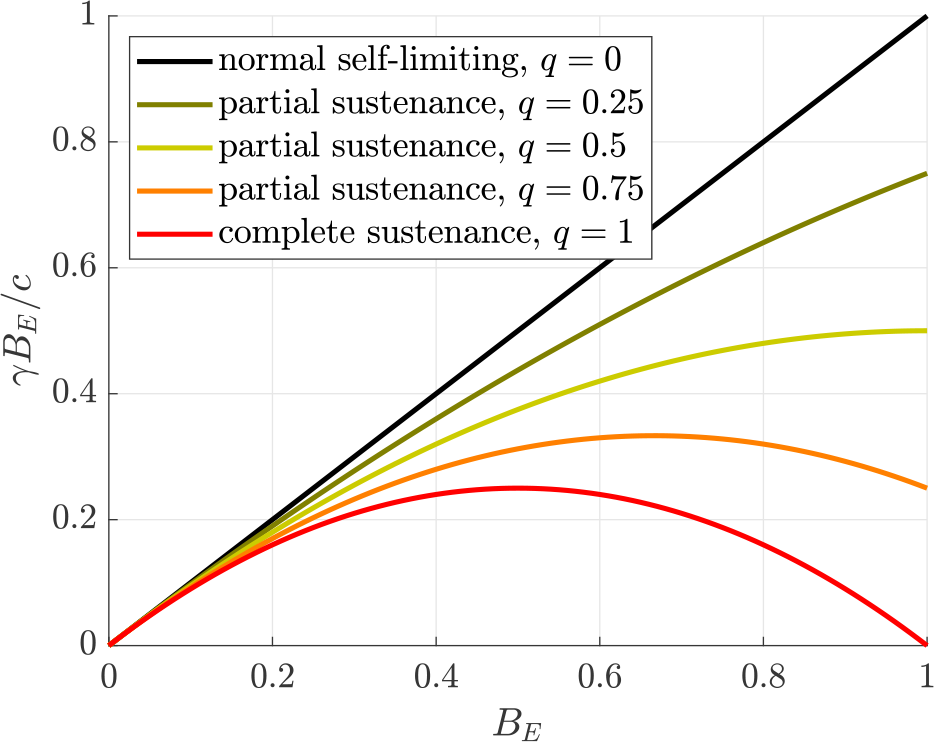
The effect of the sustenance parameter *q* on the normalized net decay *γB*_*E*_ */c*.

Including second-order effects in excitation, the non-trivial arm of the *B*_*E*_ nullcline is given by

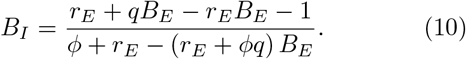

The phase portrait of the system with these second-order effects is shown in Fig. 3. The *B*_*E*_ = 0 arm of the *B*_*E*_ nullcline and the *B*_*I*_ nullcline are represented as green and blue curves respectively. The non-trivial arm of the *B*_*E*_ nullcline is plotted for different values of the sustenance parameter *q*. All other parameters are defined as previously considered in Fig. 1. In this case, increasing the value of the sustenance parameter *q* would cause the non-trivial arm of the *B*_*E*_ nullcline and consequently the fixed point corresponding to the normal state to move further northwest. This is because a higher value of *q* enables a higher proportion of excitatory bursting activity to be sustained. Further, the activity-free state remains a saddle and the normal state remains a stable spiral when the sustenance parameter is taken to be 0 *< q <* 1.

**FIG. 3.**
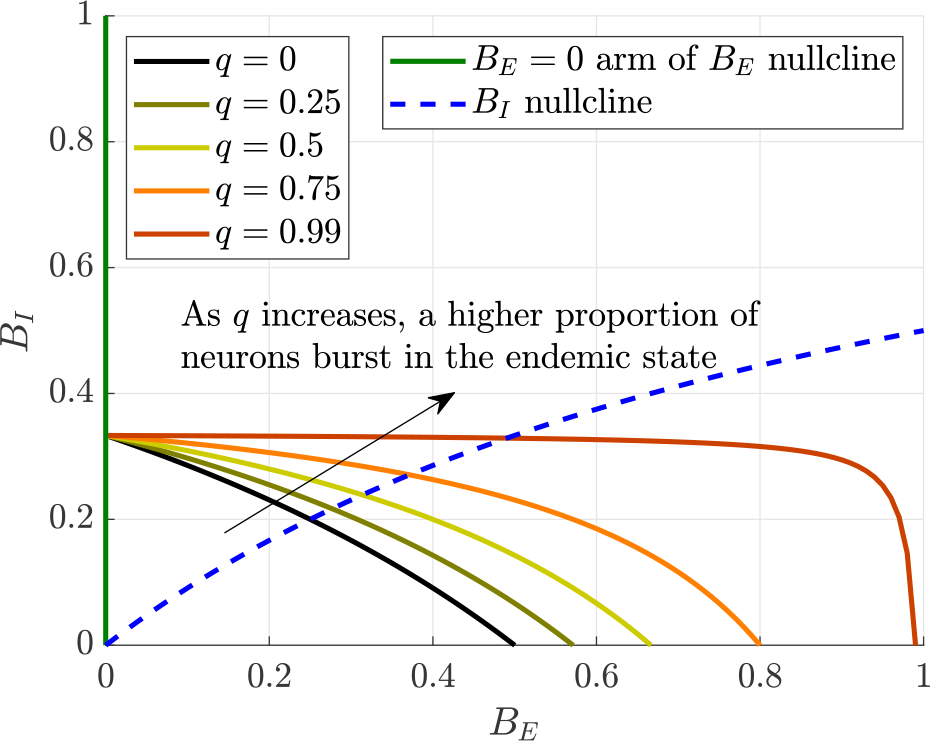
The phase portrait of the system for *β* = *b* (1 − *B*_*I*_) and *γ* = *c* (1 + *ϕB*_*I*_) (1 − *qB*_*E*_), showing the second arm of the *B*_*E*_ nullcline for different values of the sustenance parameter *q*.

In contrast to the case 0 *< q <* 1, the system behaviour alters drastically when *q* is set equal to one as seen in the next subsection.

### D. A model for seizure activity

In Section II C, we observed when *q* = 1 the sustenance perfectly counterbalanced decay at *B*_*E*_ = 1. This means that when all excitatory neurons are bursting (*B*_*E*_ = 1), they continue to burst and so net decay is zero. Further, when *B*_*E*_ = 1, there are no susceptible excitatory neurons left, and consequently, recruitment also drops to zero. Thus, setting *q* = 1 leads to 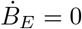 at *B*_*E*_ = 1, creating a third arm of the *B*_*E*_ nullcline. This also leads to the creation of a fixed point at 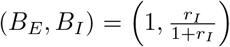. Since this is a phenomenological model, we consider this fixed point as reflecting all excitatory neurons bursting in a seizure-like state. This fixed point could either be stable or a saddle depending on the value of the parameters *r*_*E*_, *r*_*I*_, and *ϕ*.

Further, by setting *q* = 1, the non-trivial arm of the *B*_*E*_ nullcline reduces to 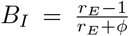, a horizontal line. The phase portrait of the system for *q* = 1, with the *B*_*E*_ and *B*_*I*_ nullclines represented as green and blue curves respectively, is shown in Fig. 4. The three arms of the *B*_*E*_ nullcline, two vertical at *B*_*E*_ = 0 and *B*_*E*_ = 1, and one horizontal at 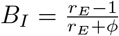 are shown in the figure.

**FIG. 4.**
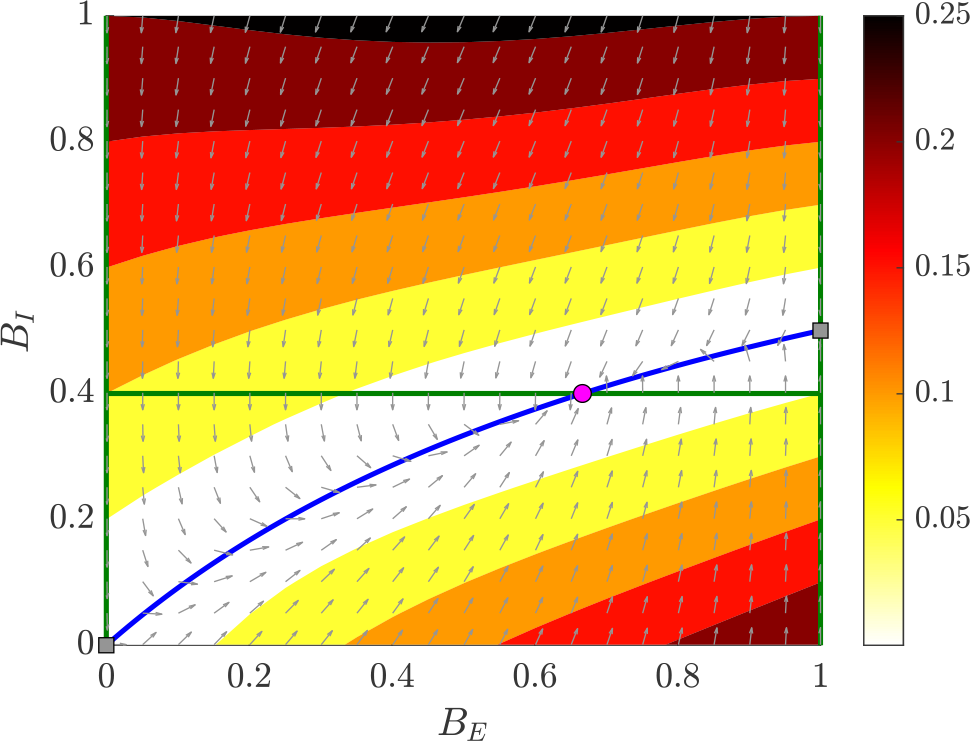
The phase portrait of the system for *β* = *b* (1 *− B*_*I*_) and *γ* = *c* (1 + *ϕB*_*I*_) (1 *− B*_*E*_), with three fixed points.

The *B*_*I*_ nullcline will always intersect with the two vertical arms of the *B*_*E*_ nullcline, giving rise to the fixed points (0, 0) and 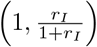 which are indicative of the activity-free state and the seizure-like state respectively. If the *B*_*I*_ nullcline intersects the horizontal arm of the *B*_*E*_ nullcline, a normal state exists and is given by 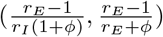. In this scenario, the activity-free state and the seizure-like state are saddles.

When the *B*_*I*_ nullcline lies entirely below the horizontal arm of the *B*_*E*_ nullcline, the normal state does not exist. This happens when

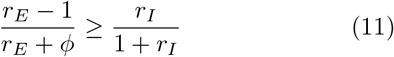

The destruction of the normal state occurs through a transcritical bifurcation when *r*_*E*_ = 1 + *r*_*I*_ (1 + *ϕ*), leading to the seizure-like state becoming stable. Simplifying Eq. (11), we find

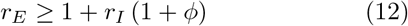

Practically, the condition given by Eq. (12) can be interpreted as follows. The model has three parameters namely, the propagation ratio for excitatory neurotransmission *r*_*E*_, that of inhibitory neurotransmission *r*_*I*_, and the scaling factor *ϕ* describing the effectiveness o f inhibition in promoting the decay of excitatory neurons into susceptibility relative to their inherent rate of decay. As such, all these parameters need to be positive to be physiologically valid.

In the absence of inhibition, the condition given by Eq. (12) reduces to *r*_*E*_ ≥ 1. Hence the necessary and sufficient condition for the seizure-like state to be stable is that the propagation ratio for excitatory neurotransmission must be greater than or equal to one.

Adding inhibition tempers the validity of the condition, *r*_*E*_ ≥ 1, resulting in it being necessary but not sufficient to ensure stability of the seizure-like state. Assuming inhibition affects o nly t he r ecruitment o f excitatory neurons and not their decay leads to *ϕ* = 0. In this case, the condition given by Eq. (12) reduces to *r*_*E*_ ≥ 1 + *r*_*I*_ . Hence the difference between the net post-synaptic excitatory activity and the net post-synaptic inhibitory activity generated by a single neuron (*r*_*E*_ − *r*_*I*_) must be greater than or equal to one for the seizure-like state to be stable. When the inhibition also accelerates the decay of excitatory neurons into susceptibility, the effect of inhibition gets scaled by a factor of 1 +*ϕ*, and the sufficient condition for the seizure-like state to be stable becomes *r*_*E*_ ≥ 1 + *r*_*I*_ (1 + *ϕ*).

In summary, we have developed an SIS-based model to describe the interaction between a single excitatory and a single inhibitory neuronal population. Described by two first-order ODEs:

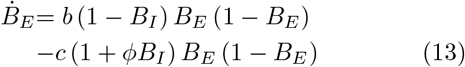

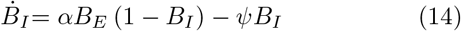

we will refer to this as the ‘Base model’ hereafter. In this model, a seizure-like state can be stabilized by choosing model parameters such that Eq. (12) is satisfied. However, such a choice implies that brain function is continuously altered such that there are no normal epochs of brain activity and frequent seizures. Such behaviour is reported in people with certain genetic encephalopathies, where important inhibitory and excitatory conductances are compromised [33].

In contrast, seizures are far more intermittent for most people with chronic epilepsy, representing less than 1% of the total brain activity [34]. Consequently, the normal state must be stable most of the time, and seizures, in this case, must arise from occasional, transient, activity-dependent imbalances between excitatory and inhibitory neurotransmission. Such imbalances can momentarily stabilize the seizure-like state even when condition Eq. (12) is not satisfied. We consider how this might occur in the next section.

## III. ROLE OF ALTERED INHIBITORY MECHANISMS IN SEIZURE GENERATION

It has long been established that epileptiform activity in the brain can be acutely induced by blocking synaptic and voltage-gated inhibitory conductances [35, 36] or by activating synaptic and voltage-gated excitatory conductances [37]. Extending this idea to the intact brain in people with epilepsy, seizures can be thought of as arising from transient, episodic, shifts in the balance between excitatory and inhibitory neurotransmission [34]. To determine why this happens, we consider what happens to the base model given in equations (13) and (14) in the absence of inhibition (*B*_*I*_ = 0):

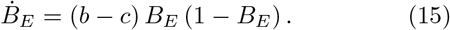

Hence, in the absence of inhibition the system has two fixed points, *B*_*E*_ = 0 and *B*_*E*_ = 1, corresponding to the activity-free and seizure-like states respectively. Further, when 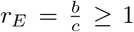, the seizure-like state will be stable. Otherwise, the activity-free state is stable. We only consider *r*_*E*_ ≥ 1 since this is necessary for the propagation of neuronal activity.

This shows that, when *r*_*E*_ ≥ 1, inhibitory neurotransmission, acting as a negative feedback, is responsible for the creation and stability of the normal state, preventing the perpetual run-off of excitatory neurotransmission into a seizure. A reduction in the negative feedback offered by inhibition or the introduction of a positive feedback could cause the seizure-like state to gain stability. This could happen in two ways: (i) the seizure-like state can gain stability, leading to coexisting normal and seizure-like attractors (known as a discontinuous or first-order transition) or (ii) the normal state could be destroyed in a saddle-node bifurcation, and the seizure-like state becomes the only stable attractor (known as a continuous or second-order transition) [38]. The proposed model is able to capture both types of transitions.

In this section, we describe two mechanisms by which the dysfunction of inhibitory neurotransmission could lead to a seizure. GABAergic interneurons are regarded as the primary inhibitory neurons and they inhibit the activity of pyramidal neurons, the primary excitatory neu-rons, through the neurotransmitter gamma-aminobutyric acid (GABA) [39]. First, we describe how inhibitory neurons inhibiting the activity of other inhibitory neurons could destroy the normal state and stabilize the seizure-like state. Subsequently, we show how an impaired release of the neurotransmitter GABA due to sustained high levels of bursting in the inhibitory population could lead to coexisting normal and seizure-like attractors.

### A. Second order effects in inhibitory neurotransmission

In Section II C, we examined second-order effects in excitatory neurotransmission and how this helps to sustain the bursting activity in the excitatory population. We will now examine the second-order effects in inhibitory neurotransmission.

Unlike excitatory neurons, the effect of bursting inhibitory neurons on each other does not promote the sustenance of activity. Instead, bursting inhibitory neurons can counteract the effect of bursting excitatory neurons and prevent the recruitment of susceptible inhibitory neurons into bursting. Furthermore, GABAergic neurotransmission due to bursting of inhibitory neurons causes postsynaptic neurons to remain in a hyperpolarized state, thereby increasing the threshold of stimulus required to fire. Hence, second-order effect of bursting inhibitory neurons would limit the activation of susceptible inhibitory neurons and promote the decay of bursting inhibitory neurons.

To capture these second-order effects we let the inhibitory recruitment and decay rates, *α* and *ψ* respectively, be functions of *B*_*I*_ . Assuming recruitment to decrease and decay to increase in a linear fashion with respect to *B*_*I*_, we choose, *α*(*B*_*I*_) = *a*(1 − *B*_*I*_) and *ψ*(*B*_*I*_) = *d*(1 + *χB*_*I*_). Here, the constants *a* and *d* denote inherent rates of recruitment and decay respectively. Consequently, the propagation ratio for the inhibitory population becomes, 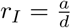.

The constant *χ* scales the second-order effects of inhibition on the decay of bursting inhibitory neurons relative to their inherent decay rate. A similar scaling term is not used to model recruitment since we require the recruitment to reach the maximum rate, *α*_max_ = *a* when *B*_*I*_ = 0, and reach zero when all the inhibitory neurons are active, *α*_min_ = 0 when *B*_*I*_ = 1.

Substituting these forms of *α* and *ψ* in Eq. (14) results in the following equation determining the evolution of the inhibitory population,

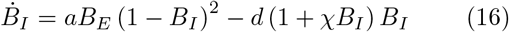

The second-order effects in inhibition do not directly affect the evolution of the excitatory population and so the corresponding equation remains the same as in the base model (Eq. (13)).

Hence, only the *B*_*I*_ nullcline is altered from that of the base model and is now defined by,

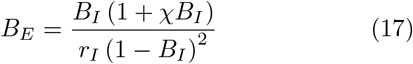

As in the base model, for the seizure-like state to be stable the *B*_*I*_ nullcline must not intersect with the horizontal arm of the *B*_*E*_ nullcline.

Now, we assess the impact of including these second-order effects in inhibition in comparison to the base model. To recall, in the base model, the *B*_*I*_ nullcline is given by (rewriting Eq. (7) to express *B*_*E*_ as a function of *B*_*I*_),

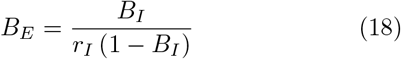

To assess these second-order effects on recruitment alone, we set *χ* = 0 in Eq. (17). The resulting expression contains an additional 1 − *B*_*I*_ factor in the denominator compared to Eq. (18). Since 1 − *B*_*I*_ ≤ 1 ∀ *B*_*I*_ ∈ [0, 1], this factor pushes the *B*_*E*_ coordinate corresponding to any *B*_*I*_ further right when compared to the base model. Overall, this leads to the *B*_*I*_ nullcline as described in Eq. (17) moving further right and further down compared to that of the base model as shown in Fig. 5. This has two implications. First, that the normal state (if stable) is pushed farther right compared to the base model. Second, the *B*_*I*_ nullcline will intersect the *B*_*E*_ = 1 arm of the *B*_*E*_ nullcline at a lower value of *B*_*I*_ compared to the base model.

**FIG. 5.**
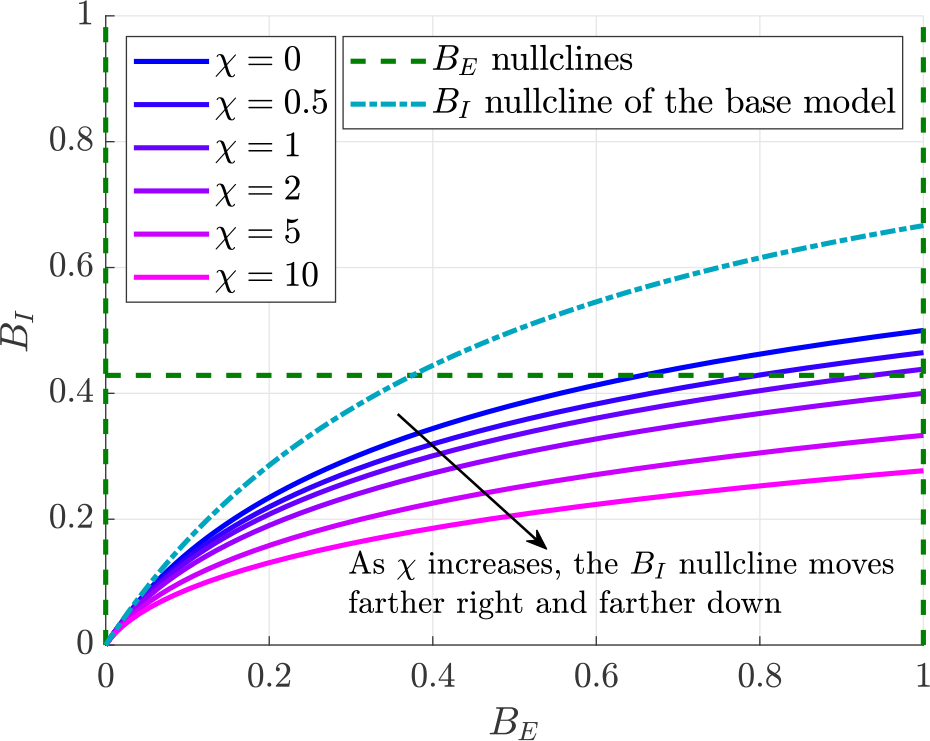
Impact of second order effects in inhibition on the *B*_*I*_ nullcline compared to the base model.

Figure 5 shows the impact of varying *χ* on all three arms of the *B*_*E*_ nullcline (dashed lines), the *B*_*I*_ nullcline of the base model (dashed and dotted lines), and the *B*_*I*_ nullcline in the presence of second-order effects in inhibition (solid lines). All nullclines being plotted by holding the parameters *r*_*E*_, *r*_*I*_ and *ϕ* constant. Note that the *B*_*E*_ nullclines are unaffected by second-order effects in inhibition and remain the same as the base model. In Fig. 5, it can be seen that the *B*_*I*_ nullcline of the base model intersects with the horizontal arm of the *B*_*E*_ nullcline, creating a normal state. The *B*_*I*_ nullcine corresponding to *χ* = 0 also intersects the horizontal arm of the *B*_*E*_ nullcline, but the normal state, in this case, is further right compared to the base model.

Both these effects are further impacted when second-order effects in decay are also considered (*χ >* 0). The presence of 1 + *χB*_*I*_ term in the numerator of Eq. (17) pushes the *B*_*I*_ nullcline further right and further down as *χ* increases as shown in Fig. 5. It could be seen that as *χ* increases from 0 to 0.5 and then to 1, the normal state moves further right.

Hence, these second-order effects serve to reduce the effect of inhibition and promote a higher proportion of excitatory neurons bursting in the normal state when it is stable. This observation finds some support in the experimental literature. For example, somatostatin-expressing GABAergic interneurons (SOM INs) that are present in layer 4 of the neocortex, preferentially inhibit fast spiking inhibitory interneurons (FS INs) in layer 4 over excitatory principal neurons (PNs). Experimentally, it has been shown that the activity of SOM INs reduces the firing of FS INs and so relieves the inhibition produced by FS INs onto PNs, and consequently enhances the output of PNs [40]. This is indicative of SOM INs providing a strong second-order effect in inhibition.

Further, the downward movement of the *B*_*I*_ nullcline implies that the *B*_*I*_ -coordinate of the point at which it would intersect the *B*_*E*_ = 1 arm of the *B*_*E*_ nullcline progressively approaches zero with increasing *χ*, depicted in Fig. 5. Hence, for the seizure-like state to be stable becomes progressively less stringent with increasing *χ* when compared to the base model. This is evident from Fig. 5, where increasing *χ* from 1 to 2 causes the *B*_*I*_ nullcline to not intersect the horizontal arm of the *B*_*E*_ nullcline. The normal state disappears in a transcritical bifurcation and the seizure-like state becomes stable. This shows that, due to second-order effects in inhibition, the seizure-like state could become stable even for the choice of parameters *r*_*E*_, *r*_*I*_ and *ϕ* that correspond to normal brain activity in the base model.

A complete derivation of stability conditions for the seizure-like state in the presence of second-order effects in inhibition, alongside an analytical proof that these conditions are less stringent than those of the base model is provided in Appendix A.

### B. Impact of impaired GABAergic neurotransmission

Seizures may also be caused due to an impairment in GABAergic neurotransmission. For example, if inhibitory neurons fire for prolonged periods of time at high frequencies, the production of GABA may be insufficient to meet the demand required by their activity. This can lead to the exhaustion of the neurotransmitter GABA at the postsynaptic excitatory neurons, reducing the level of inhibitory feedback, leading to seizures [41, 42]. This phenomenon is often referred to as ‘GABA depletion’ colloquially within the epilepsy community.

We model the GABA depletion phenomenon by introducing a quantity we term “effective inhibition”, 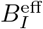 denoting the level of inhibition felt by the postsynaptic excitatory neurons. 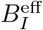 takes into account the exhaustion of the neurotransmitter GABA due to the level of activity in the inhibitory population over a time period [*t* − Δ*t, t*], and is defined by,

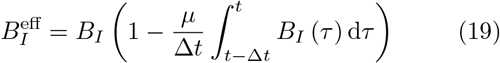

where *μ* ∈ [0, 1] is a constant that governs the level of depletion in the neurotransmitter GABA, *t* is the current time and Δ*t* is the length of the consequential time interval.

Applying Taylor’s theorem to expand the integral in Eq. (19) about Δ*t* = 0 yields,

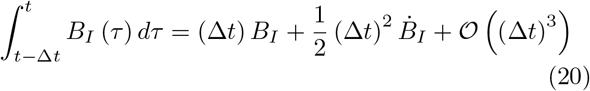

Taking the first-order approximation of the above expansion and substituting into Eq. (19) yielding a local approximation of the effective inhibition,

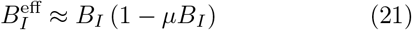

Since the effective inhibition corresponds to the level of inhibition felt by the postsynaptic excitatory neurons, we replace *B*_*I*_ with 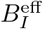 in equation Eq. (13) (which determines the evolution of the excitatory population in the base model) to account for GABA depletion,

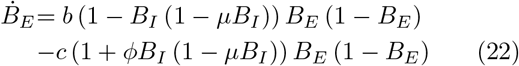

The equation determining the evolution of the inhibitory population and the *B*_*I*_ nullcline remain the same as in the base model.

The first two arms of the *B*_*E*_ nullcline are given by *B*_*E*_ = 0 and *B*_*E*_ = 1, the same as that of the base model. The remaining arms of the *B*_*E*_ nullcline differ from that of the base model and are horizontal lines defined by,

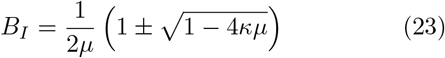

where, 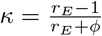

When 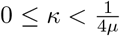, both the horizontal arms may exist. The two arms are at their extrema when *κ* = 0. As *κ* increases, the two arms come closer together. Eventually, the two arms coalesce into one at 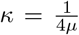 as 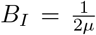.

When 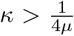, all the horizontal arms given by Eq. (23) cease to exist.

We now examine the fixed points of the system. We need to consider multiple scenarios since the *B*_*E*_ nullcline may have two to four branches dependent on the values of *κ* and *μ*. The two vertical arms of the *B*_*E*_ nullcline, *B*_*E*_ = 0 and *B*_*E*_ = 1 always intersect the *B*_*I*_ nullcline at (0, 0) and the seizure-like state 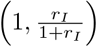, respectively. The two horizontal arms may or may not be present and if present, may or may not intersect the *B*_*I*_ nullcline. Accordingly, we present a summary of these possible scenarios in Table II.

**Table I.**
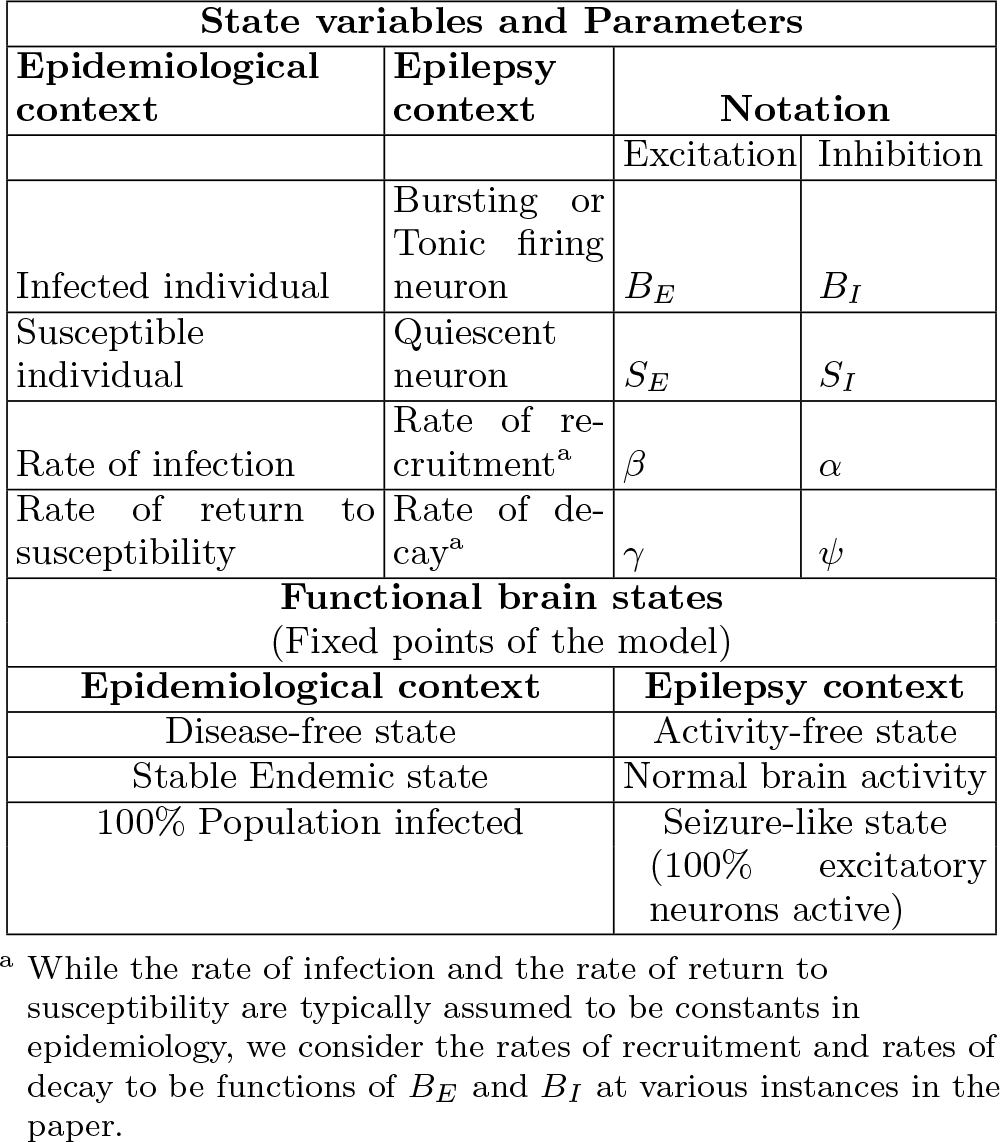
A summary of terminologies introduced in the formulation of the base model.

**Table II.** The variation in the number of fixed points and their stability for different scenarios in the GABA depletion model.

| Condition | Sub-condition | Fixed-points |
| --- | --- | --- |
| $r_E \geq r_E^c$ | | Activity-free state: Saddle<br>Seizure-like state: Stable |
| $r_E < r_E^c$ | $r_I \leq r_I^l$ | Activity-free state: Saddle<br>Seizure-like state: Stable |
| | $r_I^l < r_I \leq r_I^u$ | Activity-free state: Saddle<br>Normal state: Stable<br>Seizure-like state: Saddle |
| | $r_I > r_I^u$ | Activity-free state: Saddle<br>Normal state: Stable<br>A saddle functional brain state<br>Seizure-like state: Stable<br>(Bistability) |

In Table II, the quantities 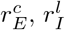 and 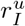 correspond to critical points at which bifurcations occur. The condition 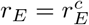 is simply the condition 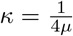 restated in terms of *r*_*E*_ and *μ*. As such, 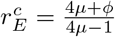. The term 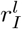 represents the critical value of *r*_*I*_ at which the *B*_*I*_ nullcline would in-tersect the lower horizontal arm of the *B*_*E*_ nullcline. This occurs when 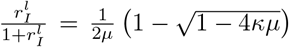 or equivalently at 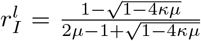. Similarly, the term 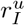 represents the critical value of *r*_*I*_ at which the *B*_*I*_ nullcline inter-sects the upper horizontal arm of the *B*_*E*_ nullcline and is given by 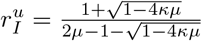.

#### 1. Scenario 1: 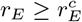

When 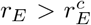, only the two vertical arms of the *B*_*E*_ nullcline exist. This results in the presence of two fixed points: the activity-free state at (0, 0) and the seizure-like state at 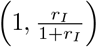. The activity-free state is always a saddle and the seizure-like state is always stable irrespective of the value of *r*_*I*_ . This contrasts significantly with the base model, where the condition for the seizure-like state to be stable depends on both *r*_*E*_ and *r*_*I*_ : *r*_*E*_ ≥ 1 + *r*_*I*_ (1 + *ϕ*).

At the critical point 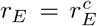, the *B*_*E*_ nullcline also includes a horizontal arm given by 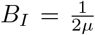. This arm exists within the domain of interest Ω only when *μ* ≥ 0.5. This horizontal arm, if present, may or may not intersect the *B*_*I*_ nullcline depending the value of *r*_*I*_ . When present, this intersection produces a saddle fixed point. Consequently, the activity-free state is always a saddle and the seizure-like state is always stable.

To understand why the seizure-like state is always stable when 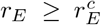 irrespective of the values taken by other parameters, we contrast the time histories of excitatory and inhibitory activity in the GABA depletion model with those of the base model in Fig. 6. For *r*_*I*_ = 0.1, the evolution of activity in the GABA depletion model is very similar to that of the base model as shown in Fig. 6 (a). In both models, since the value of *r*_*I*_ is very low, the proportion of inhibitory neurons required to keep excitatory activity “in check” is not met, and so the steady-state behaviour approaches the seizure-like state.

**FIG. 6.**
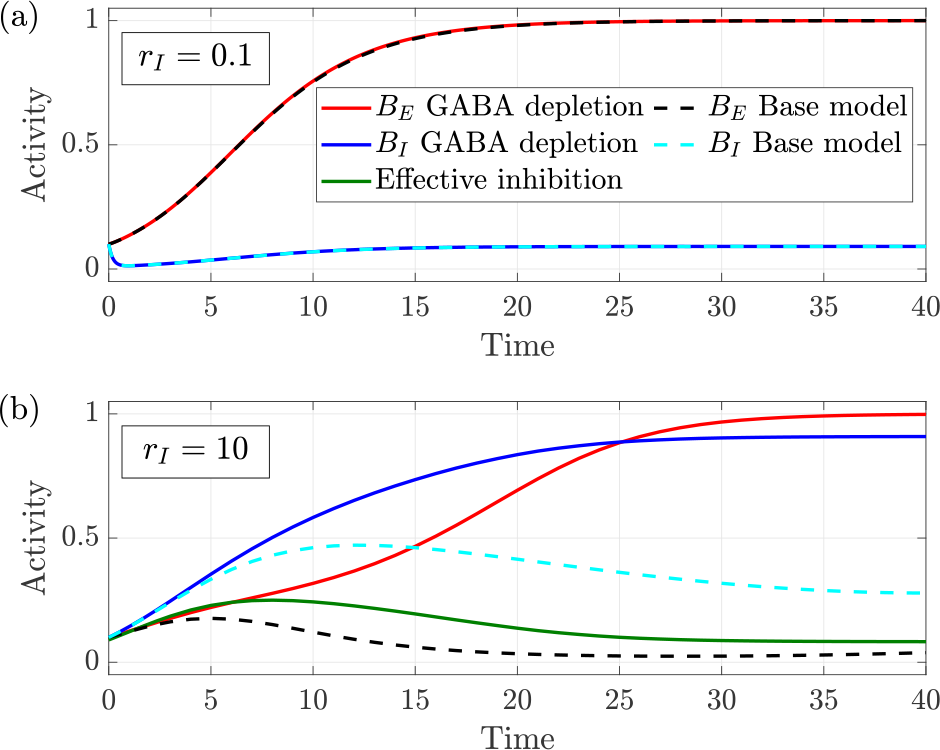
Time histories comparing the evolution of excitatory and inhibitory activity in the GABA depletion model with that of the base model for two different values of *r*_*I*_ : (a) *r*_*I*_ = 0.1 and (b) *r*_*I*_ = 10. Other parameters are fixed as follows: *r*_*E*_ = 2, *ϕ* = 1 and *μ* = 1.

On the other hand, when the value of *r*_*I*_ is very high, inhibitory neurons are recruited quickly to control excitatory activity in the base model and the steady state behaviour approaches the normal state as shown in Fig. 6 (b). However, in the GABA depletion model, this rapid recruitment of inhibitory neurons leads to GABA depletion, thereby reducing the effective inhibition. It can be seen in Fig. 6 (b) that the effective inhibition starts to fall when approximately 40% of the inhibitory neurons are recruited. The effective inhibition denotes the level of the neurotransmitter GABA available to inhibit post-synaptic excitatory neurons and as it falls, the excitatory neurons continue to burst unhindered and their steadystate behaviour approaches the seizure-like state. This shows how rapid recruitment of inhibitory neurons can lead to GABA exhaustion and inevitably to a seizure-like state.

#### 2. Scenario 2: 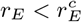

When 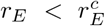, all four arms of the *B*_*E*_ nullcline (two horizontal and two vertical) may exist. The lower horizontal arm is defined by 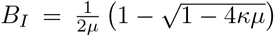 whilst the upper horizontal arm is given by 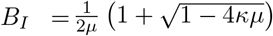

When 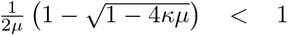, both horizontal arms vanish. In this scenario we will have only two fixed points:the activity-free state being a saddle and the seizure-like state being stable. When 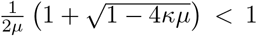, only the lower horizontal arm will exist. Here the *B*_*E*_ nullcline has two vertical arms and one horizontal arm which is qualitatively similar to the base model. If 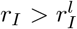, the *B*_*I*_ nullcline intersects the horizontal arm creating a normal state. In this scenario, the seizure-like state is a saddle otherwise the seizure-like state will be stable. In the specific case *μ* = 0 there is no GABA exhaustion and, consequently, the GABA depletion model converges to the base model.

When 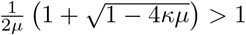, both horizontal arms of the *B*_*E*_ nullcline exist. Since this case is qualitatively different from previous cases, we consider it in more detail. The phase portrait for *μ* = 1, indicating all four arms of the *B*_*E*_ nullcline as dashed lines, is shown in Fig. 7.

Since the *B*_*I*_ nullcline is monotonically increasing for all Ω, it may intersect each of the horizontal arms of the *B*_*E*_ nullcline at most once, depending on the value of *r*_*I*_ . We consider these possible scenarios in Fig. 7.

**FIG. 7.**
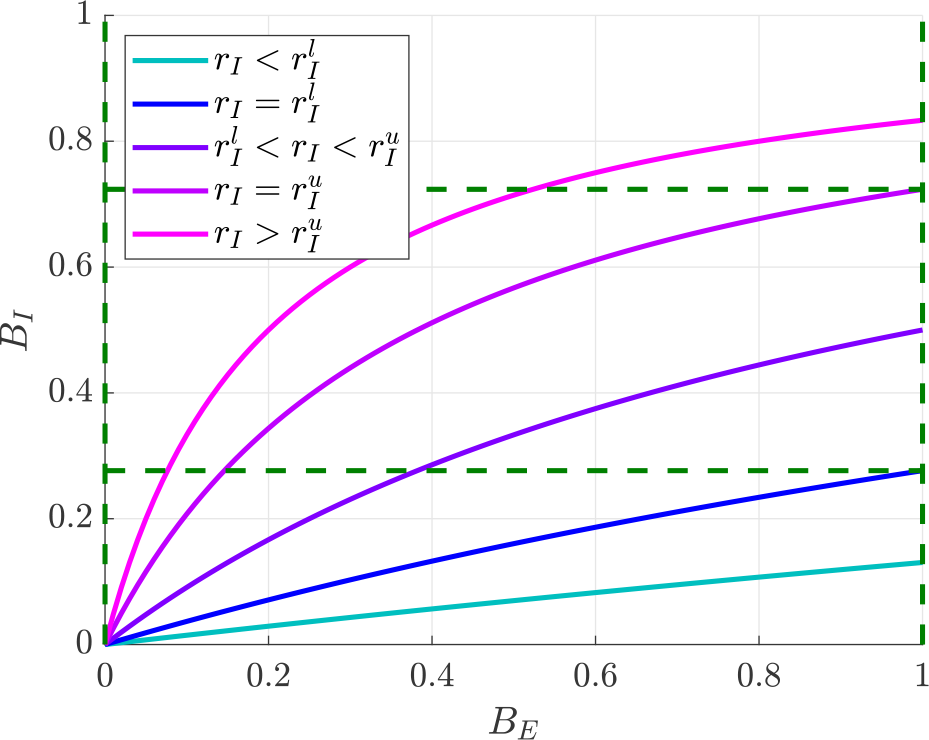
Phase portrait indicating the four arms of the *B*_*E*_ nullcline as dashed lines for the Scenario 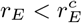 with *r*_*E*_ = 2, *ϕ* = 3 and *μ* = 1. The *B*_*I*_ nullcline is plotted as solid lines for different values of *r*_*I*_, indicating the different possibilities as to how many arms of the *B*_*E*_ nullcline would the *B*_*I*_ nullcline intersect.

If 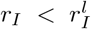, the *B*_*I*_ nullcline will not intersect both horizontal arms. Hence, we will have two fixed points, with the activity-free state being a saddle and the seizure-like state being stable.

When 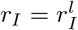, a transcritical bifurcation occurs. Consequently, when *r*_*I*_ is increased higher than 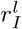, the *B*_*I*_ nullcline intersects the lower horizontal arm of the *B*_*E*_ nullcline, creating a third fixed point: a normal state and the seizure-like state loses its stability. The activity-free state remains a saddle.

When *r*_*I*_ is further increased, a further transcritical bifurcation occurs when 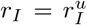. For 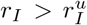, the *B*_*I*_ nullcline also intersects the upper horizontal arm of the *B*_*E*_ nullcline, creating a fourth fixed point. This fixed point is a saddle functional brain state and the seizure-like state regains stability. Effectively, we have bistability in the system: the stable normal state and the stable seizure-like state separated by a saddle functional brain state. The activity-free state remains a saddle.

This variation in the number of fixed points in the system with an increase in *r*_*I*_ is summarized in the bifurcation diagram in Fig. 8.

**FIG. 8.**
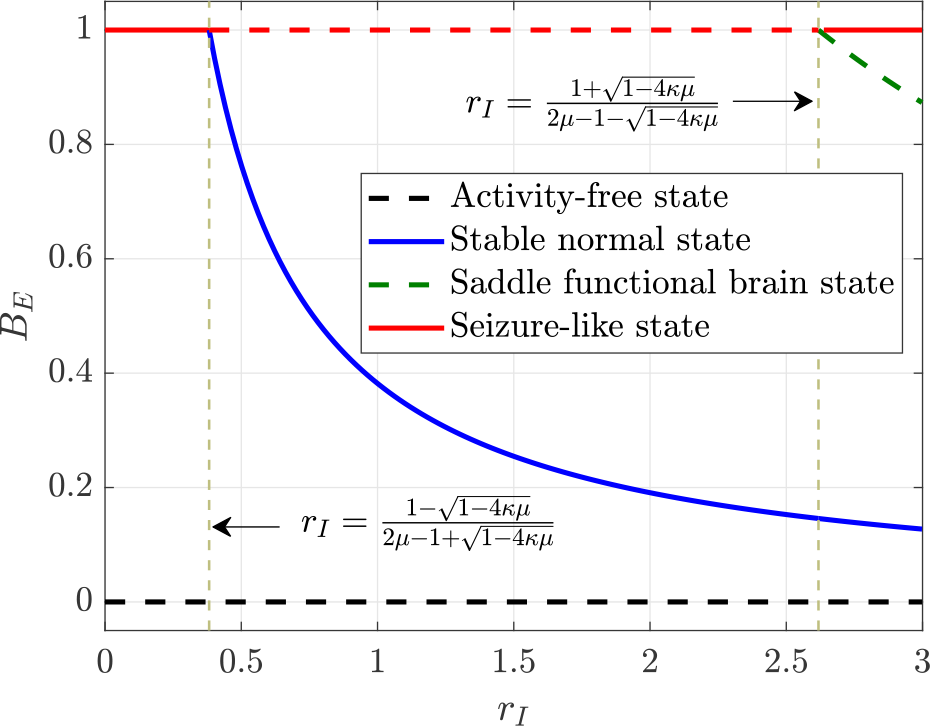
Bifurcation diagram indicating the variation in the number of fixed points with respect to *r*_*I*_ for the Scenario 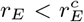 with *r*_*E*_ = 2, *ϕ* = 3 and *μ* = 1. Solid lines indicate stable fixed points while dashed lines indicate saddle fixed points.

Further, in the bistable regime, a huge increase in the activity of inhibitory neurons could cause the system to jump from the stable normal state to the seizure-like state through a first-order or discontinuous critical transition. This is because, at high levels of activity of the inhibitory population, the availability of post-synaptic GABA becomes depleted, leading to a seizure. Thus, there is an activity-dependent switch from the stable normal state to the stable seizure-like state, but this jump is possible only for a narrow range of parameters.

## IV. DISCUSSION AND OUTLOOK

In this work, we developed a model for epileptic activity that describes the interaction between a single excitatory and a single inhibitory neuronal population using a compartmental modelling approach inspired by epidemiological models. We sacrifice single-cell morphology and ion channel dynamics of a single neuron and mean-field approximation of a neuronal population, choosing instead to explore how the bursting activity propagates through interconnected neuronal populations. This enables our model to act as a platform for exploring the various mechanisms that may give rise to a seizure.

### A. Applications of our framework

Our framework would find application in the simulation of cortico-thalamic circuits and the design of experiments probing such circuits. For instance, optogenetics uses photic (i.e., light-based) stimulation to activate or inhibit a specific subset of neurons and works by incorporating genes corresponding to light-gated ion channels [43]. These light-gated ion channels, also called as opsins, allow for the efflux of ions in response to light of a specific wavelength. Thus, they act as powerful tools to turn on and off cortical circuits of interest, and have been used to explore the physiological mechanisms responsible for various neurological diseases, including epilepsy [44, 45]. Just as in optogenetic experiments, our model could also allow for the selective activation and deactivation of neuronal populations and subsequently study their effect on the overall cortical circuitry.

Epilepsy presents a difficult entity to study because of the large number of network pathologies that initiate or propagate seizures [46]. Our model could facilitate the investigation of multiple pathologies through relatively inexpensive computer simulations. Insights derived from this model can then be tested in a laboratory environment through further optogenetic experiments. Furthermore, the simulations could also be used to inform the optimal design of these optogenetic experiments, including those that involve the use of optogenetic stimulation to abort seizures [47].

### B. Limitations of the current model

A limitation of the current model is that the absolute refractory period is not modelled explicitly. Including the effects of this refractoriness would result in a two population SIRS model described by four first order ODEs. As this is the first work in applying compartmental models to epilepsy, we wanted to give a visual picture of our model. The current two population model based on the SIS framework is described by two first-order ODEs, making it possible to visualize the phase portrait of the system, including nullclines and equilibrium positions, with ease. We will incorporate the effect of the absolute refractory period in our future work.

Yet another limitation of the two population model proposed here is that the effect of increased excitation could only be captured as an increase in the parameter *r*_*E*_. Consequently, in the current form, the model cannot be used to study how a localised increase in excitatory activity and synchronization could recruit surrounding neurons and propagate further. However, this limitation could be overcome by considering multiple excitatory populations and establishing the conditions for when the neuronal activity could propagate from one population to another.

Further, our model currently captures only the proportion of neurons active at any given time and these active neurons may or may not fire in unison. As such, it cannot capture the synchronization of neurons within a population. This limitation could be addressed by assuming the phases and firing frequencies of neurons of every population to follow a certain distribution and then tracking the evolution of these distributions with time. For instance, it could be assumed that, initially, the firing frequencies follow a normal distribution and the phases follow a uniform distribution. Subsequently, the evolution of these distributions with time could be formulated as a system of stochastic differential equations and incorporated into the model. This would help us track the level of synchronization in each neuronal population with time.

### C. Future work

Epilepsy is a disease of cortical network disorganization, and as such, it is imperative to develop a model that consists of a network of multiple interconnected excitatory and inhibitory populations [48]. In the future, we will extend our model to include multiple populations and explore how the insights gathered from the single population model in this paper translate on to a network level. In particular, a key insight from our single population model is the establishment of the condition for the seizure-like state to become stable in terms of the propagation ratios of excitatory and inhibitory neurotransmission, *r*_*E*_ ≥ 1 + *r*_*I*_ (1 + *ϕ*). While the propagation ratios are taken as constants in the two population case described in this paper, it will be crucial to explore the effect of the network structure on the global equivalents of *r*_*E*_ and *r*_*I*_ when multiple populations are considered. This would help us ascertain whether a pulse of activity will percolate through the entire network or remain confined to a small number of adjacent populations, and whether such activity will quickly die out or endure indefinitely. To this end, we will also study network motifs of order two and three to distinguish the motif structures that may either enable the spread of activity from one population to another or prevent the spread and terminate the seizure.

## V. SUMMARY AND CONCLUSION

We present a novel approach describing the use of compartmental models in epilepsy to study the interaction between excitatory and inhibitory neurotransmission. We set out the base model, which contains one excitatory and one inhibitory population, and follows a two population SIS framework. This two population model is described by two first-order ODEs and has three types of equilibrium solutions, an activity-free state in which no neurons are firing, normal brain activity where a constant proportion of excitatory and inhibitory neurons are firing, and a seizure-like state in which all the excitatory neurons are firing. We then establish the condition required for stabilising a seizure-like state within the base model in terms of the propagation ratios of excitatory and inhibitory neurotransmission.

We then describe two different mechanisms by which a dysfunction in inhibitory neurotransmission could lead to seizure-like activity. When there is an impairment in GABAergic neurotransmission due to sustained high level of activity in the inhibitory population, the seizure-like state becomes co-existent with the normal state and a rapid recruitment of inhibitory neurons could lead to a jump from the normal state to the seizure-like state. When inhibitory neurons inhibit other neurons within their population, the normal state loses its stability and simultaneously, the seizure-like state becomes stable. Thus, our results further corroborate the observation of bifurcations indicative of both first-order and second-order critical transitions at the seizure onset in intracranial electroencephalographic (iEEG) signals [20].

Our approach focuses on tracking the proportion of neurons bursting within a neuronal population at any given time. Thus, our model provides a simple yet powerful framework for studying the selective activation and deactivation of neuronal populations and subsequently their effect on the overall cortical circuitry. Extended to include a network consisting of multiple neuronal populations, our model could act as a test bed for exploring the various mechanisms that may give rise to a seizure and explore the factors that enable the spread of epileptic activity through the network.

## ACKNOWLEDGMENTS

We acknowledge the support of EPSRC through the fellowship EP/T027703/1.

## Appendix A: Second order effects in inhibitory neurotransmission

In this section, we establish the conditions for the seizure-like state to be stable in the presence of second order effects in inhibitory neurotransmission and show that these conditions are less stringent than that of the base model.

For the seizure-like state to be state, the *B*_*I*_ nullcline must remain completely below the horizontal arm of the *B*_*E*_ nullcline. Since the *B*_*I*_ nullcline is monotonically increasing in Ω, the highest point in the nullcline is where it intersects the *B*_*E*_ = 1 arm of the *B*_*E*_ nullcline. This point is given by,

When *r*_*I*_ *> χ*,

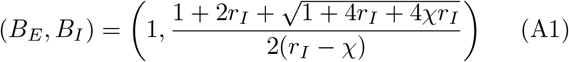

When *r*_*I*_ *< χ*,

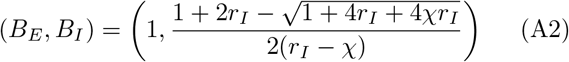

When *r*_*I*_ = *χ*,

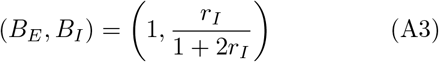

Thus, for the seizure-like state to be stable, the *B*_*I*_ - coordinate of this point must be less than or equal to 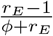. If this condition is not satisfied, the *B*_*I*_ nullcline would also intersect the horizontal arm of the *B*_*E*_ null-cline creating a stable normal state.

Further, the aforementioned condition for a seizure-like state to be stable in the presence of inhibitory-inhibitory interactions is less stringent than that of the base model and the conditions become progressively less stringent as *χ* increases. We provide an analytical proof for the same.

Let *B*^0^ and *B*^*∗*^ denote the *B*_*I*_ -coordinate of the point of intersection of the *B*_*I*_ nullcline and *B*_*E*_ = 1 arm of the *B*_*E*_ nullcline for the base model and the model with second order effects in inhibition respectively. Substituting *B*_*I*_ = *B*^*∗*^ in Eq. (17) and *B*_*I*_ = *B*^0^ in Eq. (18) and equating the RHS gives,

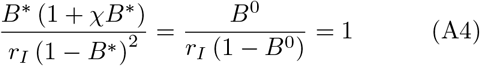

Let *B*^0^ = *B*^*∗*^ + *δ*. Substituting this in Eq. (A4),

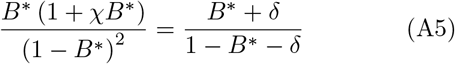

Solving for *δ* yields,

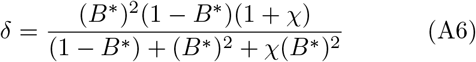

For *B*^*∗*^ ∈ (0, 1), each term enclosed in parentheses in Eq. (A6) is greater than one, implying that *δ >* 0 ∀ *χ* ≥ 0. Hence, *B*^0^ *> B*^*∗*^ ∀ *χ* ≥ 0. Thus, the condition for the seizure-like state to be stable is less stringent in the presence of second order effects in inhibition.

It should be noted that when *B*^*∗*^ = 0 or *B*^*∗*^ = 1, *δ* = 0 implying *B*^*∗*^ = *B*^0^.

To further show that it becomes progressively less stringent with an increase in *χ*, we need to show *δ* monotonically increases with an increase in *χ*. Taking the first derivative of *δ* with respect to *χ* yields,

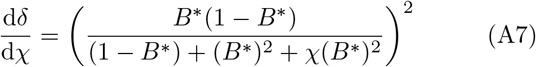

The first derivative, 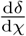, is the square of a real number as *B*^*∗*^ ∈ [0, 1] and *χ* ≥ 0. Thus, the first derivative is always greater than or equal to zero, implying that *δ* monotonically increases as a function of *χ*. Hence, the condition for the seizure-like state to be stable becomes progressively less stringent as *χ* increases.

## References

[1] P. N. Banerjee, D. Filippi, and W. A. Hauser, The descriptive epidemiology of epilepsy — a review, Epilepsy research 85, 31 (2009).

[2] S. Kalitzin, D. Velis, P. Suffczynski, J. Parra, and F. L. Da Silva, Electrical brain-stimulation paradigm for estimating the seizure onset site and the time to ictal transition in temporal lobe epilepsy, Clinical neurophysiology 116, 718 (2005).

[3] D. R. Freestone, L. Kuhlmann, D. B. Grayden, A. N. Burkitt, A. Lai, T. S. Nelson, S. Vogrin, M. Murphy, W. D’Souza, R. Badawy, et al., Electrical probing of cortical excitability in patients with epilepsy, Epilepsy & Behavior 22, S110 (2011).

[4] O. Benjamin, T. H. Fitzgerald, P. Ashwin, K. Tsaneva-Atanasova, F. Chowdhury, M. P. Richardson, and J. R. Terry, A phenomenological model of seizure initiation suggests network structure may explain seizure frequency in idiopathic generalised epilepsy, The Journal of Mathematical Neuroscience 2, 1 (2012).

[5] E. Magiorkinis, K. Sidiropoulou, and A. Diamantis, Hallmarks in the history of epilepsy: epilepsy in antiquity, Epilepsy & behavior 17, 103 (2010).

[6] M. A. Dichter and G. Ayala, Cellular mechanisms of epilepsy: a status report, Science 237, 157 (1987).

[7] J. E. Niemeyer, P. Gadamsetty, C. Chun, S. Sylvester, J. P. Lucas, H. Ma, T. H. Schwartz, and E. R. Aksay, Seizures initiate in zones of relative hyperexcitation in a zebrafish epilepsy model, Brain 145, 2347 (2022).

[8] K. P. Lillis, M. A. Kramer, J. Mertz, K. J. Staley, and J. A. White, Pyramidal cells accumulate chloride at seizure onset, Neurobiology of disease 47, 358 (2012).

[9] Z. Shiri, F. Manseau, M. Lévesque, S. Williams, and M. Avoli, Interneuron activity leads to initiation of low-voltage fast-onset seizures, Annals of Neurology 77, 541 (2015).

[10] B. Elahian, N. E. Lado, E. Mankin, S. Vangala, A. Misra, K. Moxon, I. Fried, A. Sharan, M. Yeasin, R. Staba, et al., Low-voltage fast seizures in humans begin with increased interneuron firing, Annals of neurology 84, 588 (2018).

[11] B. H. Jansen and V. G. Rit, Electroencephalogram and visual evoked potential generation in a mathematical model of coupled cortical columns, Biological cybernetics 73, 357 (1995).

[12] F. Wendling, J.-J. Bellanger, F. Bartolomei, and P. Chauvel, Relevance of nonlinear lumped-parameter models in the analysis of depth-EEG epileptic signals, Biological cybernetics 83, 367 (2000).

[13] P. Robinson, C. Rennie, and D. Rowe, Dynamics of large-scale brain activity in normal arousal states and epileptic seizures, Physical Review E 65, 041924 (2002).

[14] F. Marten, S. Rodrigues, P. Suffczynski, M. P. Richardson, and J. R. Terry, Derivation and analysis of an ordinary differential equation mean-field model for studying clinically recorded epilepsy dynamics, Physical Review E 79, 021911 (2009).

[15] F. Wendling, F. Bartolomei, J. J. Bellanger, and P. Chauvel, Epileptic fast activity can be explained by a model of impaired GABAergic dendritic inhibition, European Journal of Neuroscience 15, 1499 (2002).

[16] P. Kurbatova, F. Wendling, A. Kaminska, A. Rosati, R. Nabbout, R. Guerrini, O. Dulac, G. Pons, C. Cornu, P. Nony, et al., Dynamic changes of depolarizing GABA in a computational model of epileptogenic brain: Insight for Dravet syndrome, Experimental neurology 283, 57 (2016).

[17] J. R. Terry, O. Benjamin, and M. P. Richardson, Seizure generation: the role of nodes and networks, Epilepsia 53, e166 (2012).

[18] H. Schmidt, W. Woldman, M. Goodfellow, F. A. Chowdhury, M. Koutroumanidis, S. Jewell, M. P. Richardson, and J. R. Terry, A computational biomarker of idiopathic generalized epilepsy from resting state EEG, Epilepsia 57, e200 (2016).

[19] J. Carroll, A. Warren, and U. C. Täuber, Effects of inhibitory and excitatory neurons on the dynamics and control of avalanching neural networks, Physical Review E 99, 052407 (2019).

[20] M. L. Saggio, D. Crisp, J. M. Scott, P. Karoly, L. Kuhlmann, M. Nakatani, T. Murai, M. Dümpelmann, A. Schulze-Bonhage, A. Ikeda, et al., A taxonomy of seizure dynamotypes, Elife 9, e55632 (2020).

[21] L. Edelstein-Keshet, Mathematical Models in Biology, Classics in Applied Mathematics (Society for Industrial and Applied Mathematics, 2005).

[22] M. Wenzel, J. P. Hamm, D. S. Peterka, and R. Yuste, Acute focal seizures start as local synchronizations of neuronal ensembles, Journal of Neuroscience 39, 8562 (2019).

[23] V. Magloire, J. Cornford, A. Lieb, D. M. Kullmann, and Pavlov, KCC2 overexpression prevents the paradoxical seizure-promoting action of somatic inhibition, Nature communications 10, 1225 (2019).

[24] L. Zhu and Y. Wang, Rumor spreading model with noise interference in complex social networks, Physica A: Statistical Mechanics and its Applications 469, 750 (2017).

[25] W. Liu, X. Wu, W. Yang, X. Zhu, and S. Zhong, Modeling cyber rumor spreading over mobile social networks: A compartment approach, Applied Mathematics and Computation 343, 214 (2019).

[26] M. E. Dehshalie, M. Kabiri, and M. E. Dehshali, Stability analysis and fixed-time control of credit risk contagion, Mathematics and Computers in Simulation 190, 131 (2021).

[27] K. Zhu and L. Ying, Information source detection in the SIR model: A sample-path-based approach, IEEE/ACM Transactions on Networking 24, 408 (2014).

[28] F. Brauer, Compartmental models in epidemiology, in Mathematical Epidemiology, edited by F. Brauer, P. van den Driessche, and J. Wu (Springer Berlin Heidelberg, Berlin, Heidelberg, 2008) pp. 19–79.

[29] C. von Csefalvay, 2 - simple compartmental models: The bedrock of mathematical epidemiology, in Computational Modeling of Infectious Disease, edited by C. von Csefalvay (Academic Press, 2023) pp. 19–91.

[30] W. Gerstner and W. M. Kistler, Spiking neuron models: Single neurons, populations, plasticity (Cambridge university press, 2002).

[31] G. Deco, V. K. Jirsa, P. A. Robinson, M. Breakspear, and K. Friston, The dynamic brain: from spiking neurons to neural masses and cortical fields, PLoS computational biology 4, e1000092 (2008).

[32] C. Fraser, C. A. Donnelly, S. Cauchemez, W. P. Hanage, M. D. Van Kerkhove, T. D. Hollingsworth, J. Griffin, R. F. Baggaley, H. E. Jenkins, E. J. Lyons, et al., Pandemic potential of a strain of influenza a (h1n1): early findings, science 324, 1557 (2009).

[33] Epi4K Consortium and Epilepsy Phenome/Genome Project, De novo mutations in epileptic encephalopathies, Nature 501, 217 (2013).

[34] K. Staley, Molecular mechanisms of epilepsy, Nature neuroscience 18, 367 (2015).

[35] H. Matsumoto and C. A. Marsan, Cellular mechanisms in experimental epileptic seizures, Science 144, 193 (1964).

[36] D. A. Prince and B. J. Wilder, Control mechanisms in cortical epileptogenic foci: surround inhibition, Archives of neurology 16, 194 (1967).

[37] H. Walther, J. Lambert, R. Jones, U. Heinemann, and B. Hamon, Epileptiform activity in combined slices of the hippocampus, subiculum and entorhinal cortex during perfusion with low magnesium medium, Neuroscience letters 69, 156 (1986).

[38] C. Kuehn and C. Bick, A universal route to explosive phenomena, Science advances 7, eabe3824 (2021).

[39] H. Ye and S. Kaszuba, Inhibitory or excitatory? Optogenetic interrogation of the functional roles of gabaergic interneurons in epileptogenesis, Journal of Biomedical Science 24, 1 (2017).

[40] H. Xu, H.-Y. Jeong, R. Tremblay, and B. Rudy, Neocortical somatostatin-expressing GABAergic interneurons dis-inhibit the thalamorecipient layer 4, Neuron 77, 155 (2013).

[41] D. S.-H. Shin, W. Yu, A. Sutton, M. Calos, and P. L. Carlen, Elevated potassium elicits recurrent surges of large GABAA-receptor-mediated post-synaptic currents in hippocampal CA3 pyramidal neurons, Journal of neurophysiology 105, 1185 (2011).

[42] Z. J. Zhang, J. Koifman, D. S. Shin, H. Ye, C. M. Florez, L. Zhang, T. A. Valiante, and P. L. Carlen, Transition to seizure: ictal discharge is preceded by exhausted presynaptic GABA release in the hippocampal CA3 region, Journal of Neuroscience 32, 2499 (2012).

[43] M. Kokaia, M. Andersson, and M. Ledri, An optogenetic approach in epilepsy, Neuropharmacology 69, 89 (2013).

[44] J. T. Paz and J. R. Huguenard, Optogenetics and epilepsy: past, present and future: shedding light on seizure mechanisms and potential treatments, Epilepsy currents 15, 34 (2015).

[45] K. Lillis and K. Staley, Optogenetic dissection of ictogenesis: in search of a targeted anti-epileptic therapy, Journal of neural engineering 15, 041001 (2018).

[46] J. N. Bentley, C. Chestek, W. C. Stacey, and P. G. Patil, Optogenetics in epilepsy, Neurosurgical focus 34, E4 (2013).

[47] K. Hristova, C. Martinez-Gonzalez, T. C. Watson, N. K. Codadu, K. Hashemi, P. C. Kind, M. F. Nolan, and A. Gonzalez-Sulser, Medial septal gabaergic neurons reduce seizure duration upon optogenetic closed-loop stimulation, Brain 144, 1576 (2021).

[48] M. A. Kramer and S. S. Cash, Epilepsy as a disorder of cortical network organization, The Neuroscientist 18, 360 (2012).

